# Genome-wide association study reveals a dynamic role of common genetic variation in infant and early childhood growth

**DOI:** 10.1101/478255

**Authors:** Øyvind Helgeland, Marc Vaudel, Petur B. Juliusson, Oddgeir Lingaas Holmen, Julius Juodakis, Jonas Bacelis, Bo Jacobsson, Haakon Lindekleiv, Kristian Hveem, Rolv Terje Lie, Gun Peggy Knudsen, Camilla Stoltenberg, Per Magnus, Jørn V. Sagen, Anders Molven, Stefan Johansson, Pål R. Njølstad

## Abstract

Infant and childhood growth are dynamic processes characterized by drastic changes in fat mass and body mass index (BMI) at distinct developmental stages. To elucidate how genetic variation influences these processes, we performed the first genome-wide association study (GWAS) of BMI measurements at 12 time points from birth to eight years of age (9,286 children, 74,105 measurements) in the Norwegian Mother and Child Cohort Study (MoBa) with replication in 5,235 children (41,502 measurements). We identified five loci associated with BMI at distinct developmental stages with different patterns of association. Notably, we identified a novel transient effect in the leptin receptor (*LEPR*) locus, with no effect at birth, increasing effect on BMI in infancy, peaking at 6-12 months (rs2767486, P_6m_ = 2.0 × 10^−21^, β_6m_ = 0.16) and little effect after age five. A similar transient effect was found near the leptin gene (*LEP*), peaking at 1.5 years of age (rs10487505, P_1.5y_ = 1.3 × 10^−8^, β_1.5y_ = 0.079). Both signals are protein quantitative trait loci (pQTLs) for soluble *LEPR* and *LEP* in plasma in adults and independent from signals associated with other adult traits mapped to the respective genes, suggesting novel key roles of common variation in the leptin signaling pathway for healthy infant growth. Hence, our longitudinal analysis uncovers a complex and dynamic influence of common variation on BMI during infant and early childhood growth, dominated by the *LEP*-*LEPR* axis in infancy.

---

BMI patterns in infancy and childhood follow well-characterized trajectories: a rapid increase soon after birth until approximately nine months, the adiposity peak, followed by a gradual decline until approximately 4-6 years of age and then the adiposity rebound, when BMI starts to increase again until the end of puberty^1^. Recently, a study revealed that the most powerful predictor of obesity in adolescence is an increase in BMI between two and six years of age^2^, but the underlying cause for this remains unknown. To explore how common genetic variation influences these processes, we performed the first genome-wide association study (GWAS) of early growth in the population-based birth cohort, the Norwegian Mother and Child Cohort Study (MoBa)^3^ (Supplementary Table 1). A total of 17,474 children in MoBa were genotyped in discovery and replication combined. The children’s BMI was measured at birth, 6 weeks, 3, 6, 8 months, and 1, 1.5, 2, 3, 5, 7, and 8 years of age (Fig. 1a). We performed genotype quality control (QC), imputation using the Haplotype Reference Consortium (HRC), and phenotype QC, leaving 9,286 and 5,235 samples for the discovery and replication cohorts, respectively, all of Norwegian ancestry.

**Fig. 1.**
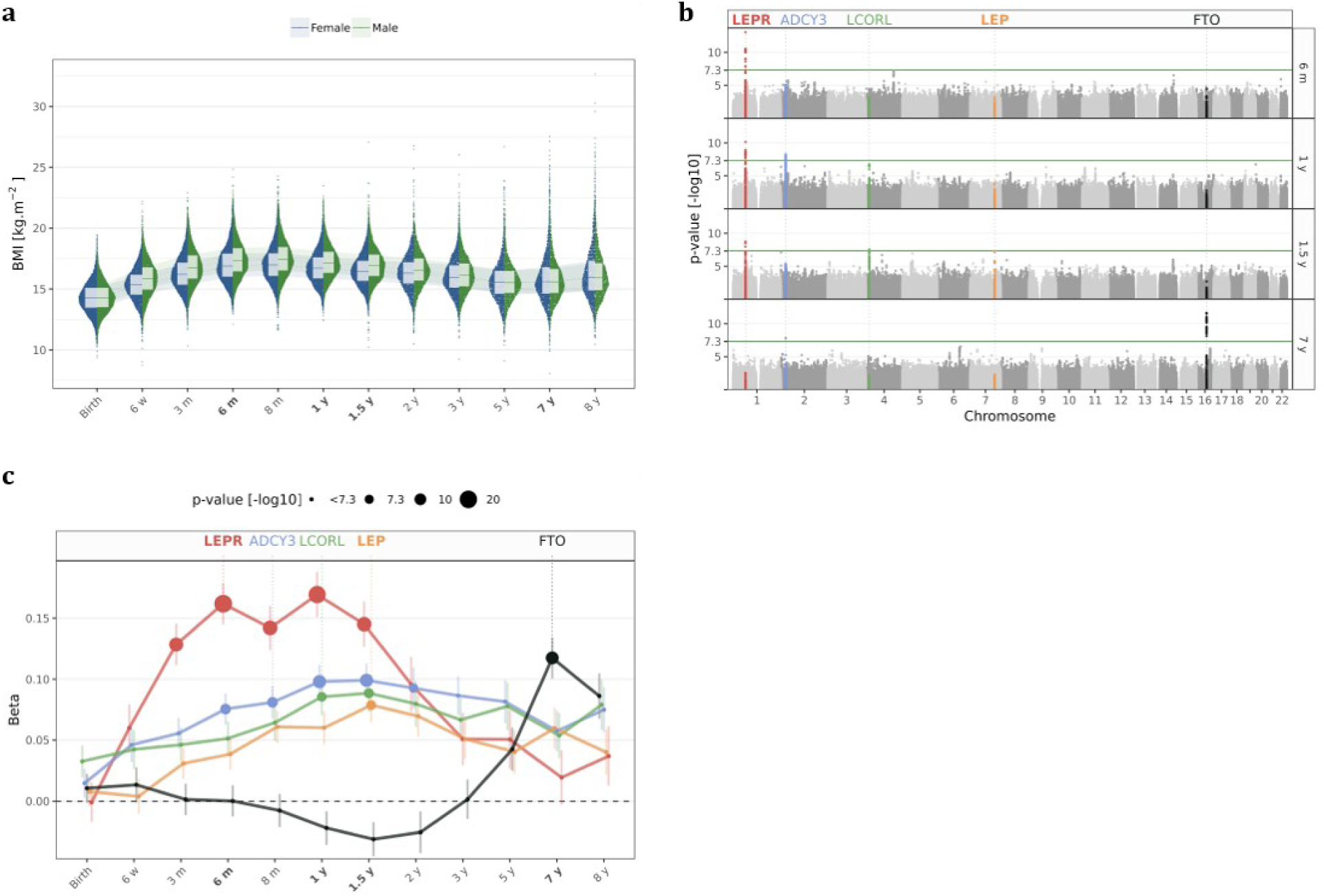
GWAS of BMI from 12 timepoints in the Mother Child Cohort of Norway. **a**, BMI values in kg/m^2^ for all samples used for association are plotted at each time point, with the normalized density of points restricting the jitter along the x-axis to the left for females (blue) and to the right for males (green). Ribbons and box plots, showing the median and quartiles, are plotted in background and foreground, respectively. **b**, Manhattan plots showing association results for the discovery sample (N=9,286) at six time points: 6 months, 1 year, 1.5 year, and 7 years. *LEPR*, *ADCY3*, *LCORL*, *LEP*, and *FTO* loci are highlighted in red, blue, green, orange, and black, respectively. **c**, Regression betas are plotted in sd units at each time point for rs2767486, rs13035244, rs6842303, rs10487505, and rs9922708, the lead SNPs of the *LEPR*, *ADCY3*, *LCORL*, *LEP*, and *FTO* loci, respectively, in the same colors as 1b. Results are presented for the meta-analysis of discovery and replication sample. The size of the points is proportional to - log10(P-value of the association). Error bars represent 1 standard error of mean (SEM) on each side of the point.

**Table 1.**
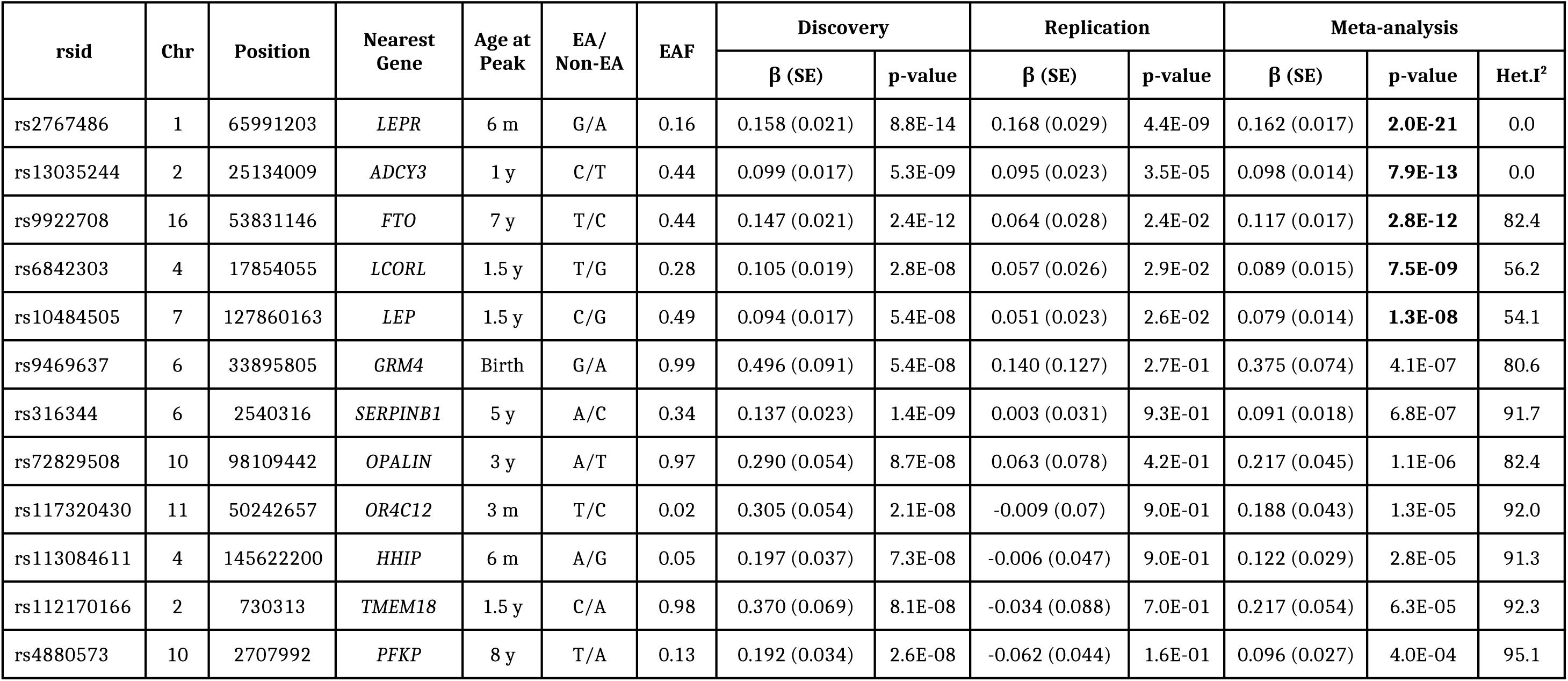
Summary statics for the signals that met criteria for replication. For each SNP, the table lists (i) the rsid; (ii) the genomic coordinates in build GRCh37; (iii) the nearest gene; (iv) the age at peak, *i.e.* lowest P-value; (v) the BMI-increasing and non-increasing alleles, EA and Non-EA, respectively; (vi) the BMI-increasing allele frequency (EAF); (vii) the regression beta (β), standard error (SE), and associated P-value for discovery, replication, and meta-analysis; and (viii) heterogeneity I^2^.

**Fig. 2.**
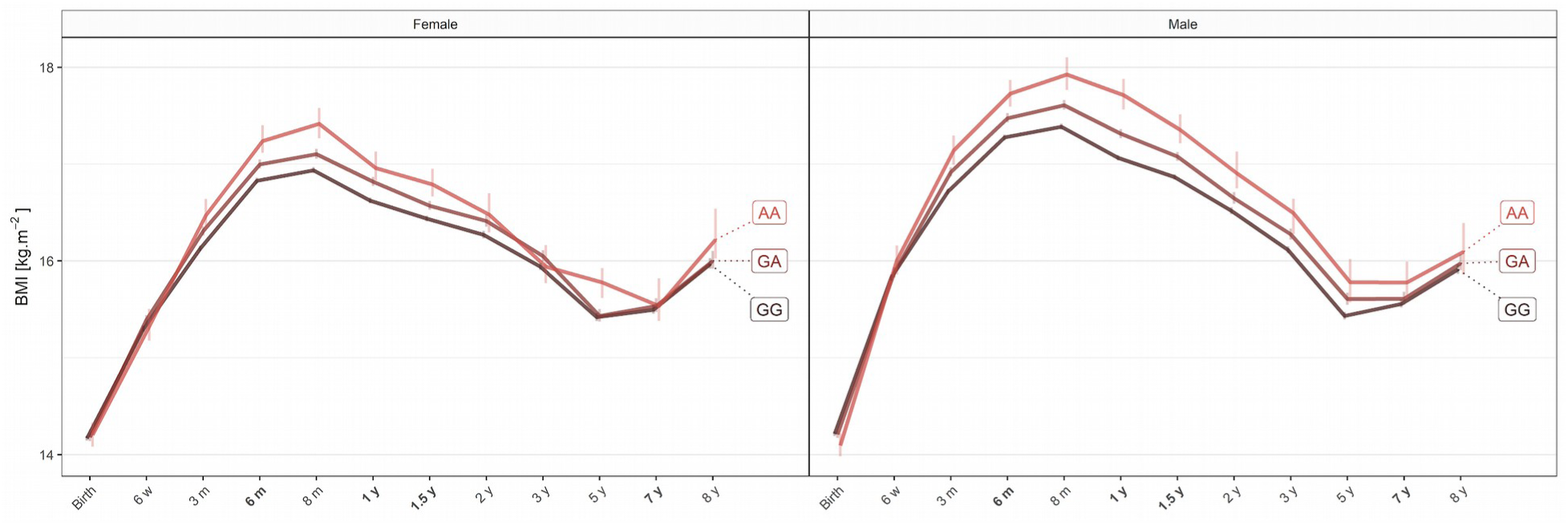
BMI trajectories stratified by genotype of the lead SNP of the *LEPR* locus (rs2767486). The median of the BMI in kg/m^2^ at each time point for all samples (discovery + replication) stratified by genotype GG, GA, or AA in black, brown, and red, respectively, for females (left) and males (right). Error bars represent 1 SEM on each side of the point. Trajectories for all five GWA-significant SNPs identified in the study are depicted in Supplementary Figure 6.

We conducted separate linear regression analyses of standardized BMI for each time point using an additive genetic model. The lead SNPs at independent loci reaching P < 10^−7^ at one or more time points in the discovery sample were taken forward for replication (Table 1). This revealed a highly dynamic pattern of association during early growth. SNPs in five independent loci reached genome-wide significance, presenting peak association at different time points: (1) a novel, intronic SNP rs2767486 in the *LEPR* locus peaking at six months; (2) an intronic SNP, rs13035244, near *ADCY3* peaking at one year; (3) an intronic SNP rs6842303 near *LCORL* peaking at 1.5 year; (4) an intergenic SNP rs10487505 near *LEP* peaking at 1.5 year; and (5) an intronic SNP rs9922708 near *FTO* peaking at seven years (Fig. 1b, c, and Supplementary Fig. 1, 2).

The strongest association with BMI was found for rs2767486 at six months (P_6m_ = 2.0 × 10^−21^, β_6m_ = 0.16) in the *LEPR*/*LEPROT* locus. The locus strongly associated with BMI from three months of age, with effects peaking at 6-12 months, and waning from age three with little effect at eight years (Fig. 1c, 2, and Supplementary Fig. 3a). We found no evidence of association at birth for rs2767486 or nearby markers in our data or in recent large publicly available GWASs of birth weight^4^ and adult BMI^5,6^. Thus, this locus most likely affects BMI development primarily during infancy. Conditioning on rs2767486 revealed a putative additional signal in the *LEPR* locus, rs17127815 (P_6m_ = 7.5 × 10^−5^ after conditioning), that mirrored the association pattern of the main signal (Supplementary Fig. 3b).

*LEPR* encodes the leptin receptor, which functions as a receptor for the adipose cell-specific hormone leptin. High leptin levels suppress hunger by interacting with the long form of the leptin receptor (OB-RL) in the hypothalamus^7^. The soluble form of leptin receptor (sOB-R), which is produced through ectodomain shedding of OB-RL in peripheral tissues, can bind leptin in circulation, and thereby reduce its effect on the central nervous system^8^. The *LEPR* locus has previously been implicated in monogenic morbid obesity^9,10^, severe childhood obesity^11^, age of menarche^12^, age of voice breaking^13^, levels of fibrinogen^14^ and C-reactive protein^15^, several blood cell count traits^16,17^, and plasma sOB-R levels^17,18^. To test whether any of the established variants for these traits explains the observed association with BMI in infancy, we repeated the analysis conditioning on the top SNPs reported in these studies. The association with infant BMI remained unaffected by conditioning on these SNPs, except for rs2767485 (Supplementary Fig. 4a), the strongest pQTL for sOB-R-plasma levels in adults^18^. Intriguingly, this SNP is located only 12.2 kbp upstream our top SNP rs2767486, with strong LD (r^2^ = 0.9) between the BMI-raising and the sOB-R-increasing alleles.

The strong association between variants in the *LEPR* locus and infant BMI suggests an important role of leptin signaling in early growth. The genome-wide significant association with infant BMI for rs10487505 located 20 kbp upstream of *LEP* is therefore noteworthy. This SNP is a known pQTL for circulating leptin levels in adults^19^. The leptin-increasing allele from Kilpeläinen et al.^19^ is associated with lower infant BMI in our data. The effect presents a rise-and-fall pattern, rising during the 3-12 months period when the *LEPR* signal is at its plateau, reaching its peak at 1.5 years (P_1.5y_ = 1.3 × 10^−8^, β_1.5y_ = 0.08) before waning (Fig. 1c, Supplementary Fig. 5c, 6). Children homozygous for the alleles associating with higher sOB-R and lower leptin levels exhibited higher mean standardized BMI (+0.65) than children homozygous for the opposite alleles (Fig. 3).

**Fig. 3.**
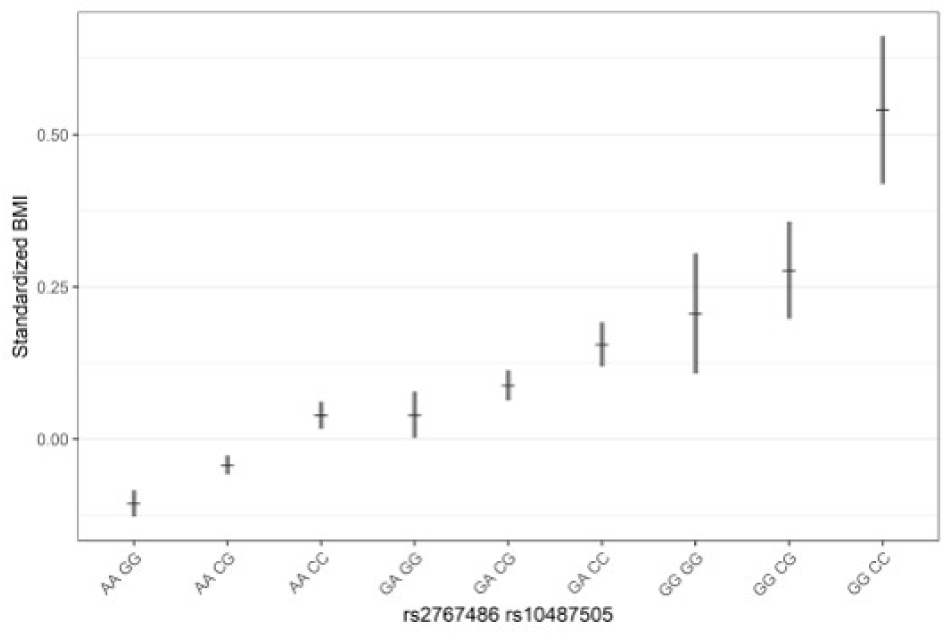
Standardized BMI at 1.5 years of the lead SNPs in the *LEPR* and *LEP* loci stratified by the combined genotypes of rs2767486 and rs10487505, respectively. The mean of the standardized BMI of all samples (discovery + replication) at 1.5 years is plotted after stratifying by genotype. Error bars represent 1 SEM on each side of the point.

We identified a novel association with BMI in the *LCORL* locus for rs6842303, presenting a similar rise-and-fall pattern with peak effect at 1.5 years (P_1.5y_ = 7.5 × 10^−9^, β_1.5y_ = 0.09) (Fig. 1c, Supplementary Fig. 5b, 6). Previously, this marker has been associated with related traits such as birth weight, birth length, infant length, and adult height. Interestingly, rs6842303 has also been associated with peak height velocity in infancy^20^, but no association was reported in the largest adult BMI GWASs to date^5,6^. This supports our finding of a transient effect of *LCORL* in early growth.

The second strongest signal was found at the *ADCY3* locus. Biallelic mutations in *ADCY3* have recently been found to cause severe syndromic obesity^21,22^. *ADCY3* is known to interact with *MC4R*, and rare mutations in *MC4R* account for 3-5% of severe obesity^23^. The lead *ADCY3* SNP, rs13035244, showed no association at birth, gradually became genome-wide significant with a peak effect between one and 1.5 years (P_1y_ = 7.9 × 10^−13^, β_1y_ = 0.10), and then stabilized during the course of childhood (Fig. 1c, Supplementary Fig. 5a, 6). This result is in agreement with a previous study of growth trajectories in children from one to 17 years of age^24^.

In contrast to the rise-and-fall pattern reported here for signals in the *LEPR*, *ADCY3*, *LEP*, and *LCORL* loci, the *FTO* risk allele displayed a trend towards a slightly negative effect around adiposity peak before gradually turning positive from three years of age, reaching genome-wide significance at seven years (P_7y_ = 2.8 × 10^−12^, β_7y_ = 0.12), in agreement with Sovio et al.^25^. (Fig. 1c, Supplementary Fig. 5d, 6)

Previous studies have suggested a tight genetic overlap between child and adult BMI, but the details of this relationship across the first years of life remain elusive^24,26^. We used LD score regression^27^ in LD Hub^28^ to quantify the shared genetic contribution between BMI at each of the 12 time points and other traits (Fig. 4a,b and Supplementary Fig. 7). Our data show that although adult BMI and other adult obesity traits normally associated with poor metabolic control were positively correlated with childhood BMI from age 5-8 years, this correlation was much weaker below the age of three years. Notably, the genetic correlation with a range of non-anthropometric traits varied substantially at infant age (Supplementary Fig. 8). Polygenic risk score analyses across all time points for markers associated with birth weight^4^, childhood BMI^26^, and adult BMI^5,6^ revealed similar patterns (Fig. 4c). We also used LD score regression to estimate the SNP-based heritability of BMI measurements across infancy and childhood. The LD score regression-based heritability estimates varied with age, with relatively modest levels at birth and during the adiposity rebound, and high levels when adiposity is high, *i.e.* around adiposity peak and from seven years of age onwards (Fig. 4a). This finding is supported by twin-studies that also show high heritability estimates for BMI in infancy, lower levels around four years of age, followed by higher estimates in later childhood^29^. Collectively, these results further indicate that the genetic mechanisms underlying BMI change from infancy to adulthood.

**Fig. 4.**
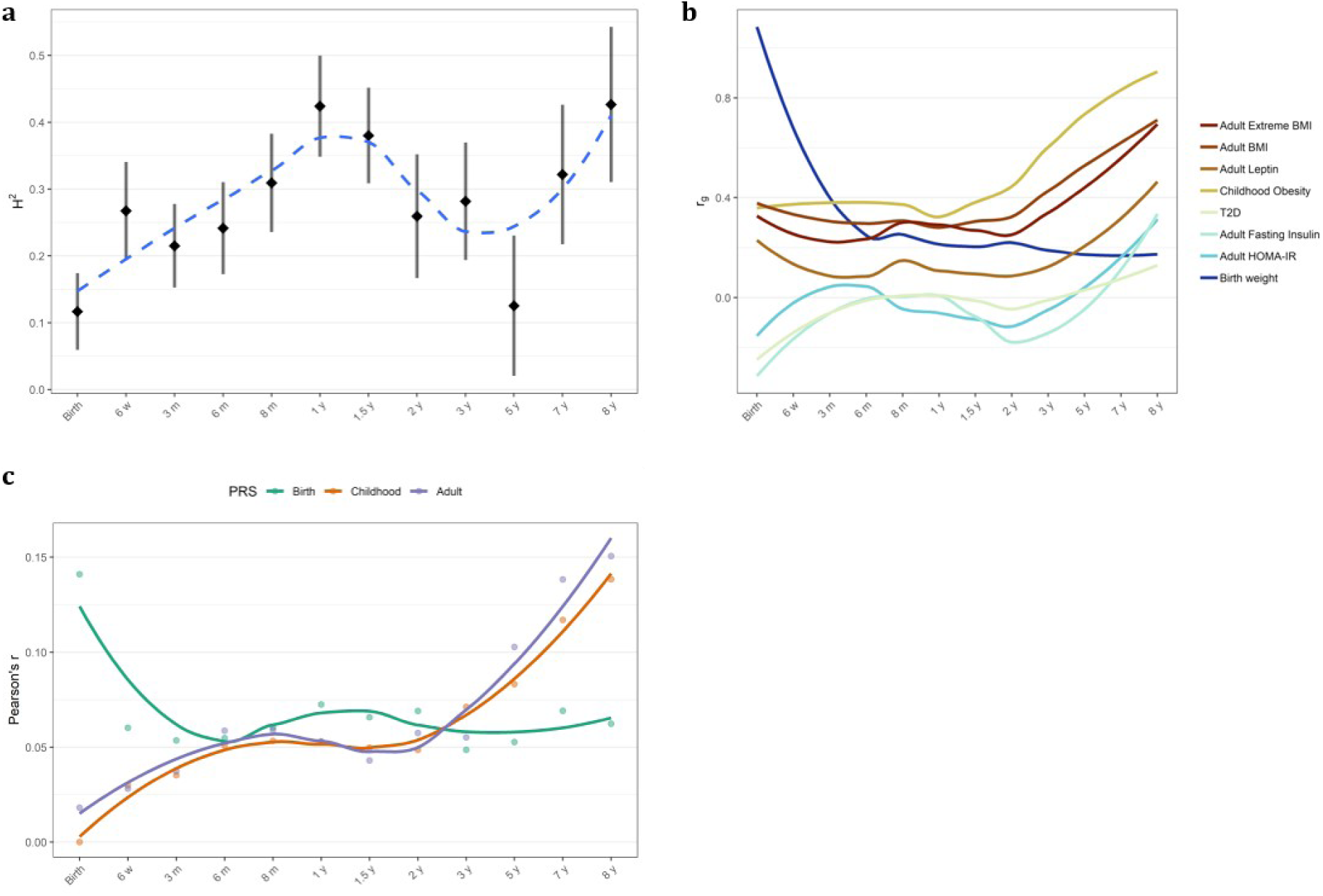
Evolution of heritability, correlation with other traits, and risk scores at all time points. **a.** H^2^ estimates based on LD score regression for BMI are plotted at each age (black) along with locally estimated scatterplot smoothing (LOESS) local regression (in blue). Error bars represent 1 SEM on each side of the point. **b**, LOESS local regression of the LD score regression coefficient (r_g_) between our BMI association results at each age and phenotypes from LD Hub (i) adult extreme BMI, (ii) adult BMI, (iii) leptin not adjusted for BMI, (iv) childhood obesity, (v) type 2 diabetes, T2D, (vi) fasting insulin main effect, (vii) homeostatic model assessment for insulin resistance, HOMA-IR. **c**, Pearson’s r of the correlation between polygenic risk scores (PRS) and BMI at each age along with LOESS local regression. Correlation with PRS for birth weight, childhood BMI, and adult BMI are displayed in green, orange, and purple, respectively.

To our knowledge, we report the first GWAS with dense measurements of BMI during the first year of life. The few GWASs published on BMI in infancy and childhood mainly involve children above five years of age, *i.e.* during adiposity rebound^24,26^. These studies point toward a strong genetic correlation for BMI around adiposity rebound and adulthood. Our results confirm a strong overlap of the genetics of BMI from five to eight years and adulthood, however, this association is much less pronounced during infancy. Infant weight and height have considerable heritable components^30^. Our results suggest that there are distinct molecular mechanisms that dynamically and specifically influence weight gain in infancy, partly acting through leptin signaling.

Leptin has an important role in fetal growth, and is positively correlated with birth weight^31^. Leptin levels are high at birth and decrease quickly, whereas sOB-R levels are low at birth and increase rapidly during the first postnatal days^32^. This pattern is hypothesized to be an important mechanism for suppressing leptin-induced energy expenditure during the first neonatal days. The sOB-R level remains very high during the first two years of life and then declines^33^, mirroring the association of *LEPR* with infant BMI observed in our study (Fig. 1c). An effect of genetic variant(s) on the level of sOB-R in infancy is therefore a possible causal mechanism underlying the association with BMI. An interaction between the *LEPR*- and *LEP*-associated variants with increased BMI in individuals who carry both the sOB-R-raising and leptin-lowering alleles would further support a mechanism where sOB-R in circulation sequesters leptin, reducing its membrane receptor activation, hence promoting energy intake during infancy. The SNPs associated with increased BMI during infancy near *LEPR* and *LEP* are not known to affect adult BMI. In fact, they are not in LD with any marker associated with adult diseases, and might thus promote healthy weight gain during infancy, a notion further supported at the genome level by LD score regression. This result is further supported by a recent independent study^34^ suggesting that SNPs in the *LEPR*/*LEPROT* locus are associated with BMI at the adiposity peak.

In summary, in our first, large GWAS performed in the Mother Child Cohort of Norway capitalizing on a wealth of phenotypes, the longitudinal analysis uncovers a complex and dynamic influence of common genetic variation on BMI during infant and early childhood growth, dominated by the *LEP*-*LEPR* axis in infancy. Improved understanding of infant weight biology is important as childhood obesity as well as undernutrition and premature births are worldwide challenges. Our study provides novel knowledge of time-resolved genetic determinants for infant and early childhood growth, suggesting that weight management intervention should be tailored to developmental stage and genetic profile of the patients. For instance, homeostatic increase in the level of sOB-R during infancy might have a positive effect on weight gain without being associated with adult overweight, offering a potential drug target for ensuring weight gain in infant care.

## Methods

### Study population

The Norwegian Mother and Child Cohort Study (MoBa) is an open-ended cohort study that recruited pregnant women in Norway from 1999 to 2008. Approximately 114,000 children, 95,000 mothers, and 75,000 fathers of predominantly Norwegian ancestry were enrolled in the study from 50 hospitals all across Norway^3^. Anthropometric measurements of the children were carried out at hospitals (at birth) and during routine visits by trained nurses at 6 weeks, 3, 6, 8 months, and 1, 1.5, 2, 3, 5, 7, and 8 years of age. Parents later transcribed these measurements to questionnaires. In 2012, the project Better Health By Harvesting Biobanks (HARVEST) randomly selected 11,490 umbilical cord blood DNA samples from MoBa’s biobank for genotyping, excluding samples matching any of the following criteria: (1) stillborn, (2) deceased, (3) twins, (4) non-existing Medical Birth Registry (MBR) data, (5) missing anthropometric measurements at birth in MBR, (6) pregnancies where the mother did not answer the first questionnaire (as a proxy for higher fallout rate), and (7) missing parental DNA samples. In 2016, HARVEST randomly selected a second set of samples, 5,984, using the same criteria.

### Genotyping

For the discovery sample, genotyping was performed using Illumina’s HumanCoreExome-12 v.1.1 and HumanCoreExome-24 v.1.0 arrays for 6,938 and 4,552 samples, respectively, at the Genomics Core Facility located at the Norwegian University of Science and Technology, Trondheim, Norway. The replication sample was genotyped using Illumina’s Global Screening Array v.1.0 for all 5,984 samples at the Erasmus University Medical Center in Rotterdam, Netherlands. We used the Genome Reference Consortium Human Build 37 (GRCh37) reference genome for all annotations and included autosomal markers only for this study.

Genotypes were called in Illumina Genome Studio (v.2011.1 for discovery and v.2.0.3 for replication). Cluster positions were identified from samples with call rate ≥ 0.98 and GenCall score ≥ 0.15. We excluded variants with low call rates, signal intensity, quality scores, heterozygote excess and deviation from Hardy-Weinberg equilibrium (HWE) based on the following QC parameters: call rate < 98%, cluster separation < 0.4, 10% GC-score < 0.3, AA T Dev > 0.025, HWE P-value < 10^−6^. Samples were excluded based on call rate < 98% and heterozygosity excess > 4 SD. Study participants with non-Norwegian ancestry were excluded after merging with samples from the HapMap project (ver. 3). Sample pairs with PI_HAT > 0.1 in identical-by-descent (IBD) calculations were resolved by removing a random sample in each pair. After genotype calling and QC, 9,286 (80.8%) from the discovery sample set, and 5,235 (87.5%) from the replication sample remained eligible for analysis.

### Pre-phasing and imputation

Prior to imputation, insertions and deletions were removed to make the dataset congruent with Haplotype Reference Consortium (HRC) v.1.1 imputation panel using HRC Imputation preparation tool by Will Rayner version 4.2.5 (see URLs): insertions and deletions were excluded. Allele, marker position, and strand orientation were updated to match the reference panel. A total of 384,855 and 568,275 markers remained eligible for phasing and imputation for the discovery and replication set, respectively. Pre-phasing was conducted locally using Shapeit v2.790^35^. Imputation was performed at the Sanger Imputation Server (see URLs) with positional Burrows-Wheeler transform^36^ and HRC version 1.1 as reference panel.

### Phenotypes

Age, height, and weight values were extracted from Medical Birth Registry (MBR) of Norway for birth, and from the study questionnaires for remaining time points. Pregnancy duration in days was extracted from MBR and pregnancies with duration < 37 × 7 days were excluded (515 pregnancies). Height and weight values were inspected at each age and those provided in centimeter or gram instead of meter and kilogram, respectively, were converted. Extreme outliers, typically an error in handwritten text parsing or a consequence of incorrect units, were excluded (47 length and 8 weight measurements). A value *x* was considered as extreme outlier if *x* >*m*+ 2*×* (*perc*_99_ *−m*) or *x* <*m−* 2*×* (*m− p erc*_1_), where *m* represents the median and *perc*_1_ and *perc*_99_ the 1^st^ and 99^th^ percentiles, respectively.

Subsequently, height and weight curves were inspected for extreme outliers by monitoring the variation of height and weight over time as follows: (i) the height and weight ratio between consecutive ages were calculated at each time point but the last: *r_i_*=*x_i+1_* / *x_i_* where *r_i_* is the ratio at time point *i* and *x_i_* is height or weight at *i*; (ii) the ratios were scaled after logarithm base 2 transformation, 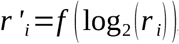, using the function *f* of equation 1:

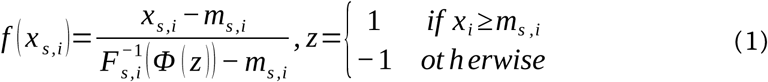

Where *x_s,i_* is the value for an individual of sex *s* at time point *i*, *m_s, i_* is the median, 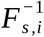 the empirical quantile function of the values at *i* of individuals of sex *s* presenting at least three values before age two (exclusive) and at least two values after age two (inclusive), and *Φ* the distribution function of the standard normal distribution; (iii) the height or weight of an individual at time point *i*, presenting surrounding scaled ratios *r*′ _i−1_ and *r*′ _i_ was considered as outlier and excluded if *r*′ _i−1_>1 and *r*′ _i_ ←1 or if *r*′ _i−1_ *←*1 and *r*′ _i_ >1, corresponding to peaks or gaps in the curve, respectively.

If for an individual of sex *s*, two consecutive height values, *h_i_* and *h_i+1_* presented a decrease in height, *i.e. h_i_* +1< *h_i_*, this was considered an artefact and corrected as follows.

If the individual presented three or more other height measurements, *h _j_* with *j ≠ i* and *j ≠ i*+1, for each *j* the corresponding height at *i* and *i* +1 was estimated by interpolating the height curve using the ratios as in equation 2:

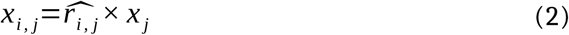

Where *x_i, j_* is the value at *i* interpolated from *j*, *x* _j_ is the value at *j*, and 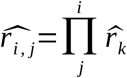 and 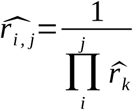, with 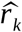 the median of the ratios *r* at time point *k* for the individuals of sex *s* presenting at least three values before age two (exclusive) and at least two values after age two (inclusive). If, for all *j*, *h_i_* >*h_i, j_*, *h_i_* was considered as outlier and excluded. Similarly, if, for all *j*, *h_i_* +1< *h_i, j_*, *h_i_* +1 was considered as outlier and excluded.

Alternatively, if the individual presented two or fewer other height measurements, and *h_i_* >*h_high_*, *h_i_* was considered as outlier and removed, with *h_high_* defined as in equation 3:

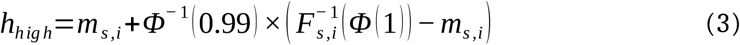

Where *m_s, i_* is the median and 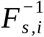 the empirical quantile function of the heights at *i* of individuals of sex *s* presenting at least three values before age two (exclusive) and at least two values after age two (inclusive), *Φ* and _*Φ*_^−1^ the distribution and quantile functions of the standard normal distribution, respectively. Similarly, if the individual presented two or less other height measurements, and *h_i_* +1< *h_low_*, *h_i_* +1 was considered as outlier and removed, with *h_low_* defined as in equation 4:

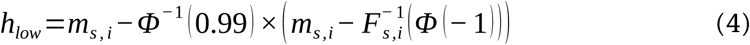

If *h_i_* and *h_i_* +1 were not considered as outliers, *h_i_* and *h_i_* +1 were defined as the median of *h_i, j_* as defined in equation 2, for all *j ≠ i* and *j ≠ i*+1, respectively. Starting from 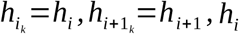 and *h_i+1_* were iteratively decreased or increased, respectively, until *h_i_* +1 *≥h_i_* as described in equation 5 and 6.

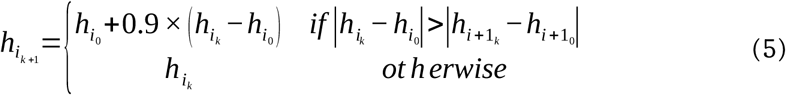

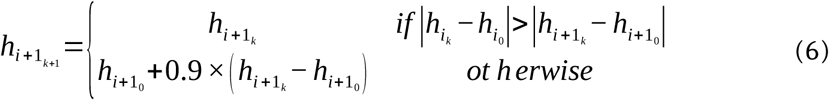

Subsequently, height and weight missing values were imputed from the individual height and weight curves at all ages for individuals presenting at least three values before age two (exclusive) and at least two values after age two (inclusive), and until age two (exclusive), for individuals presenting at least three values before age two (exclusive). A missing value at *i* was imputed to *x_i_*=*median* (*x_i, j_*), with *x_i, j_* as defined in equation 2. Importantly, missing values were imputed only if at least two non-imputed values were present at both earlier and later ages. Upon imputation of missing values, outlier removal and height decrease correction was conducted as described previously, and the new missing values were imputed using the same rules.

Finally, the body mass index (BMI) was computed where both height and weight values were available. At each time point, BMI values were scaled prior to association as described in equation 1. These scaled values are referred to as *standardized BMI* in the text.

The analysis of the phenotypes was conducted in R version 3.5.1 (2018-07-02) -- “Feather Spray” (https://www.R-project.org).

### Statistical analyses

Genome-wide analyses were performed using SNPTEST v.2.5.2 using dosages of alternate allele with an additive linear model using sex, batch, and ten principal components as covariates. LD score regression was performed with LD Hub v.1.9.0 using LDSC v.1.0.0^27^ using all markers remaining after performing pruning recommended by the LD Hub^28^ authors.

### Figures

All figures in the manuscript were generated in R version 3.5.1 (2018-07-02) -- “Feather Spray” (https://www.R-project.org). In addition to the system packages, the following packages were used: ggplot2 version 3.0.0, scico version 1.0.0, gtable version 0.2.0, ggrepel version 0.8.0, and ggdendro version 0.1-20.

### URLs

HRC or 1000G Imputation preparation and checking: http://www.well.ox.ac.uk/~wrayner/tools; Sanger Imputation Service, https://imputation.sanger.ac.uk;

## Ethics

The study was approved by the Regional Committee for Medical and Health Research Ethics in Norway (#2012/67).

## Supporting information

## Author contributions

Ø.H. and M.V. performed the analyses.

O.L., T.Z., J.J, J.B., B.J., H.L., K.H., R.T.L., G.P.K., C.S., and P.M.M. contributed to sample acquisition and genotyping.

J.J. and J.B. assisted with genotype quality control.

Ø.H., M.V., S.J., and P.R.N. wrote the manuscript with contributions from all authors.

P.B.J, J.V.S., and A.M. critically revised the manuscript for important intellectual content.

S.J. and P.R.N. designed and directed the study.

P.R.N. secured funding and initiated the study.

## Acknowledgements

This work was supported by grants (to P.R.N.) from the European Research Council (AdG #293574), the Bergen Research Foundation (“Utilizing the Mother and Child Cohort and the Medical Birth Registry for Better Health”), Stiftelsen Kristian Gerhard Jebsen (Translational Medical Center), the University of Bergen, the Research Council of Norway (FRIPRO grant #240413), the Western Norway Regional Health Authority (Strategic Fund “Personalized Medicine for Children and Adults”), and the Norwegian Diabetes Foundation; and (to S.J.) Helse Vest’s Open Research Grant. This work was partly supported by the Research Council of Norway through its Centres of Excellence funding scheme (#262700), Better Health by Harvesting Biobanks (#229624) and The Swedish Research Council, Stockholm, Sweden (2015-02559), The Research Council of Norway, Oslo, Norway (FRIMEDBIO ES547711, March of Dimes (#21-FY16- 121). The Norwegian Mother and Child Cohort Study is supported by the Norwegian Ministry of Health and Care Services and the Ministry of Education and Research, NIH/NIEHS (contract no N01-ES-75558), NIH/NINDS (grant no.1 UO1 NS 047537-01 and grant no.2 UO1 NS 047537-06A1). We are grateful to all the families in Norway who are taking part in this ongoing cohort study.

