## Supplementary material for "Genome-wide association study reveals a dynamic role of common genetic variation in infant and early childhood growth"

| Age | Discovery | | | Replication | | | Merged | | |
|---|---|---|---|---|---|---|---|---|---|
| | # Samples | Sex | BMI [Median (IQR, Q1 - Q3)] | # Samples | Sex | BMI [Median (IQR, Q1 - Q3)] | # Samples | Sex | BMI [Median (IQR, Q1 - Q3)] |
| Birth | 9,286 | F: 48.2%<br>M: 51.8% | F: 14.24 (1.67, 13.45 - 15.12)<br>M: 14.28 (1.63, 13.49 - 15.12) | 5,235 | F: 50.4%<br>M: 49.6% | F: 14.19 (1.65, 13.34 - 14.99)<br>M: 14.24 (1.67, 13.42 - 15.09) | 14,521 | F: 49.0%<br>M: 51.0% | F: 14.23 (1.68, 13.40 - 15.08)<br>M: 14.26 (1.66, 13.46 - 15.12) |
| 6 w | 6,450 | F: 47.5%<br>M: 52.5% | F: 15.36 (1.69, 14.51 - 16.20)<br>M: 15.87 (1.87, 14.93 - 16.81) | 3,444 | F: 50.3%<br>M: 49.7% | F: 15.30 (1.84, 14.38 - 16.22)<br>M: 15.83 (1.85, 14.94 - 16.79) | 9,894 | F: 48.5%<br>M: 51.5% | F: 15.34 (1.74, 14.48 - 16.21)<br>M: 15.86 (1.87, 14.94 - 16.80) |
| 3 m | 7,970 | F: 48.0%<br>M: 52.0% | F: 16.21 (1.86, 15.33 - 17.20)<br>M: 16.76 (1.94, 15.84 - 17.78) | 4,453 | F: 50.2%<br>M: 49.8% | F: 16.15 (1.94, 15.26 - 17.20)<br>M: 16.77 (1.97, 15.85 - 17.82) | 12,424 | F: 48.8%<br>M: 51.2% | F: 16.19 (1.90, 15.30 - 17.20)<br>M: 16.77 (1.95, 15.84 - 17.79) |
| 6 m | 8,017 | F: 48.2%<br>M: 51.8% | F: 16.89 (1.93, 16.00 - 17.93)<br>M: 17.35 (1.94, 16.44 - 18.37) | 4,516 | F: 50.4%<br>M: 49.6% | F: 16.90 (1.91, 15.99 - 17.91)<br>M: 17.31 (2.01, 16.37 - 18.38) | 12,531 | F: 49.0%<br>M: 51.0% | F: 16.90 (1.93, 16.00 - 17.93)<br>M: 17.34 (1.96, 16.41 - 18.38) |
| 8 m | 7,197 | F: 47.9%<br>M: 52.1% | F: 16.99 (1.89, 16.06 - 17.95)<br>M: 17.44 (1.96, 16.51 - 18.47) | 3,966 | F: 50.4%<br>M: 49.6% | F: 17.04 (1.91, 16.09 – 18.00)<br>M: 17.48 (2.08, 16.46 - 18.54) | 11,161 | F: 48.8%<br>M: 51.2% | F: 17.01 (1.89, 16.06 - 17.96)<br>M: 17.45 (2.00, 16.49 - 18.49) |
| 1 y | 6,970 | F: 48.0%<br>M: 52.0% | F: 16.71 (1.77, 15.85 - 17.62)<br>M: 17.14 (1.77, 16.31 - 18.08) | 3,839 | F: 50.1%<br>M: 49.9% | F: 16.70 (1.79, 15.87 - 17.66)<br>M: 17.17 (1.92, 16.19 - 18.11) | 10,808 | F: 48.7%<br>M: 51.3% | F: 16.70 (1.78, 15.85 - 17.64)<br>M: 17.15 (1.81, 16.28 - 18.09) |
| 1.5 y | 6,800 | F: 47.9%<br>M: 52.1% | F: 16.47 (1.70, 15.65 - 17.35)<br>M: 16.94 (1.79, 16.05 - 17.85) | 3,780 | F: 50.0%<br>M: 50.0% | F: 16.53 (1.83, 15.68 - 17.51)<br>M: 16.93 (1.81, 16.03 - 17.85) | 10,578 | F: 48.6%<br>M: 51.4% | F: 16.50 (1.76, 15.66 - 17.42)<br>M: 16.93 (1.80, 16.04 - 17.85) |
| 2 y | 4,631 | F: 48.6%<br>M: 51.4% | F: 16.31 (1.80, 15.43 - 17.23)<br>M: 16.54 (1.86, 15.67 - 17.53) | 2,502 | F: 49.4%<br>M: 50.6% | F: 16.33 (1.95, 15.35 - 17.31)<br>M: 16.66 (1.85, 15.70 - 17.55) | 7,131 | F: 48.9%<br>M: 51.1% | F: 16.32 (1.87, 15.40 - 17.27)<br>M: 16.57 (1.85, 15.68 - 17.53) |
| 3 y | 5,069 | F: 47.5%<br>M: 52.5% | F: 15.95 (1.88, 15.09 - 16.98)<br>M: 16.14 (1.93, 15.20 - 17.13) | 2,834 | F: 49.9%<br>M: 50.1% | F: 16.00 (1.98, 15.00 - 16.98)<br>M: 16.17 (1.94, 15.28 - 17.22) | 7,905 | F: 48.3%<br>M: 51.7% | F: 15.96 (1.93, 15.05 - 16.98)<br>M: 16.15 (1.91, 15.25 - 17.16) |
| 5 y | 4,245 | F: 47.7%<br>M: 52.3% | F: 15.54 (1.95, 14.58 - 16.53)<br>M: 15.52 (1.87, 14.65 - 16.52) | 2,208 | F: 51.0%<br>M: 49.0% | F: 15.39 (1.88, 14.46 - 16.35)<br>M: 15.43 (1.77, 14.57 - 16.35) | 6,454 | F: 48.8%<br>M: 51.2% | F: 15.43 (1.95, 14.51 - 16.47)<br>M: 15.49 (1.86, 14.61 - 16.47) |
| 7 y | 4,629 | F: 47.4%<br>M: 52.6% | F: 15.61 (2.23, 14.60 - 16.83)<br>M: 15.59 (2.03, 14.65 - 16.68) | 2,638 | F: 50.3%<br>M: 49.7% | F: 15.45 (2.25, 14.46 - 16.72)<br>M: 15.50 (2.00, 14.59 - 16.58) | 7,268 | F: 48.5%<br>M: 51.5% | F: 15.53 (2.23, 14.57 - 16.80)<br>M: 15.56 (2.01, 14.63 - 16.65) |
| 8 y | 3,862 | F: 47.9%<br>M: 52.1% | F: 15.98 (2.51, 14.85 - 17.36)<br>M: 15.92 (2.28, 14.88 - 17.16) | 2,087 | F: 50.3%<br>M: 49.7% | F: 15.98 (2.55, 14.80 - 17.34)<br>M: 15.87 (2.15, 14.86 - 17.01) | 5,949 | F: 48.7%<br>M: 51.3% | F: 15.98 (2.55, 14.81 - 17.36)<br>M: 15.91 (2.24, 14.88 - 17.12) |

***Supplementary Table 1***

**Summary statistics on the phenotypes used for association.** At each time point, the number of samples, share of females (F) and males (M), and BMI summary statistics are listed. The BMI summary statistics provided are Median (IQR, Q1-Q3), with IQR the inter-quartile range, and Q1 and Q3 the first and third quartiles, respectively. Summary statistics are provided for (i) the discovery cohort, (ii) the replication cohort, and (iii) both cohorts merged. Note that in the latter case, outlier detection is conducted on the entire cohort, yielding minute differences between the number of samples in the merged cohort compared to the sum of samples in discovery and replication.

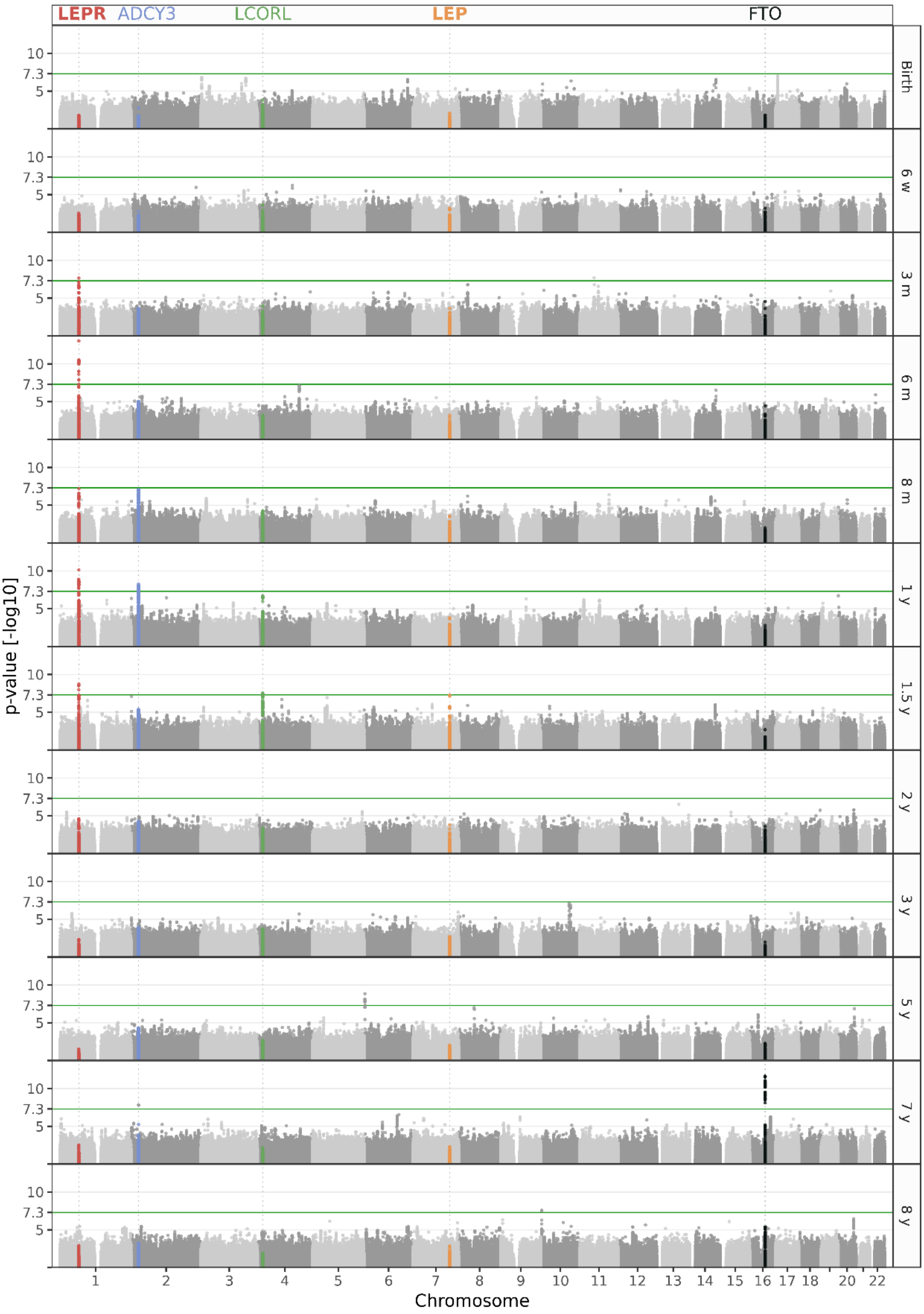

***Supplementary Figure 1***

**Manhattan plots all time points.** Manhattan plots showing association results at birth, 6 weeks, 3, 6, 8 months, and 1, 1.5, 2, 3, 5, 7, and 8 years of age in the discovery sample. *LEPR*, *ADCY3*, *LCORL*, *LEP*, and *FTO* loci are highlighted in red, blue, green, orange, and black, respectively.

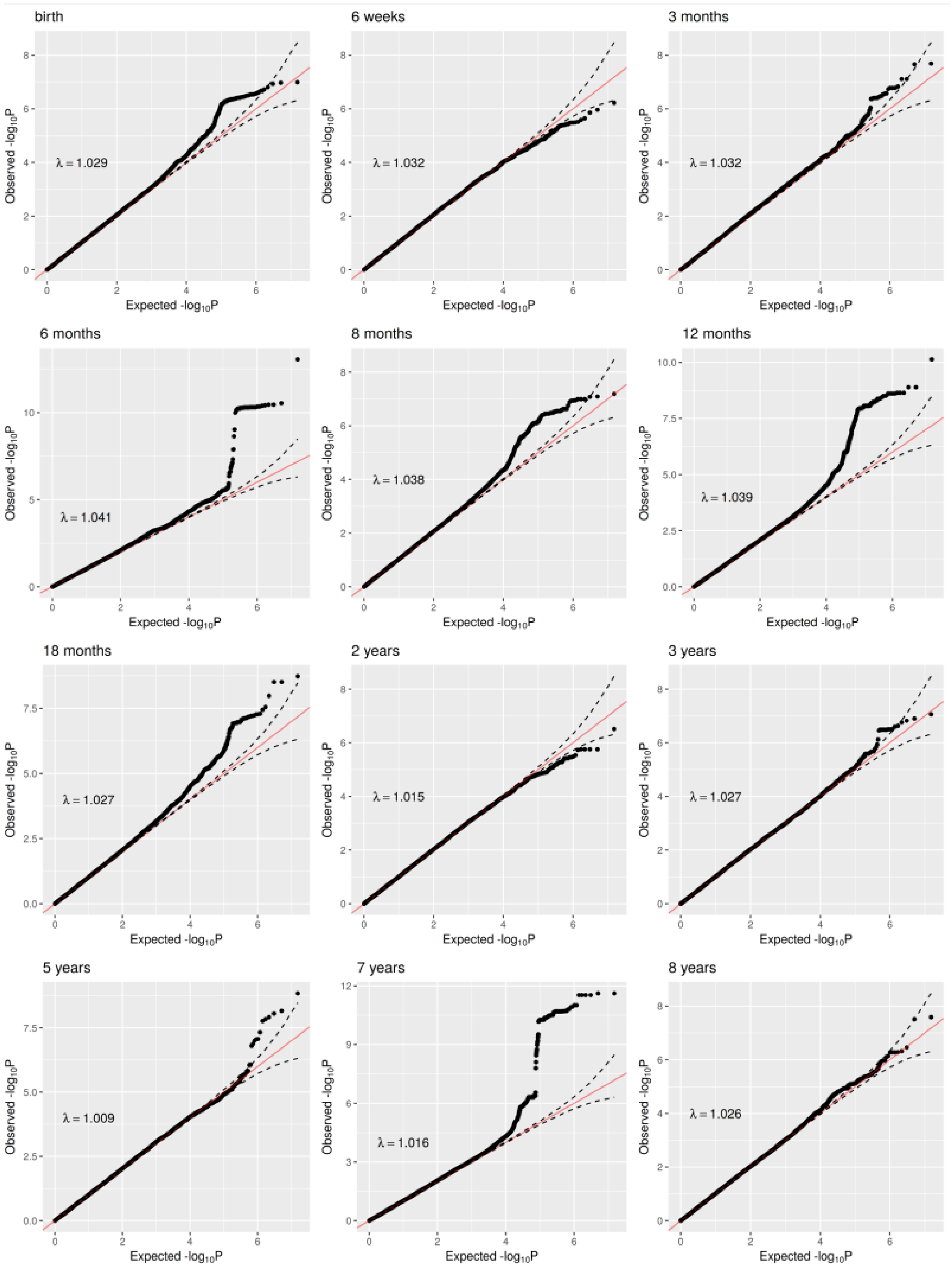

***Supplementary Figure 2***

**QQ plots all time points.** QQ plots showing association results at birth, 6 weeks, 3, 6, 8 months, and 1, 1.5, 2, 3, 5, 7, and 8 years of age in the discovery sample. λ representing the inflation at median is indicated on the plot. The diagonal (y=x) is displayed in red and 95% confidence intervals as dashed black lines.

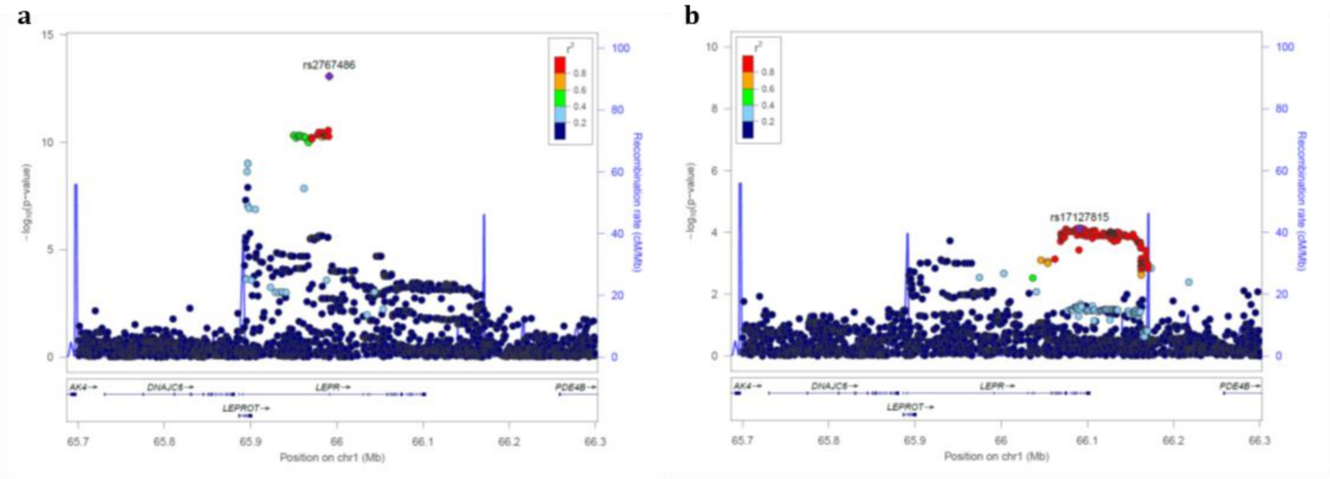

***Supplementary Figure 3***

**Novel association in the *LEPR* locus.** Regional association plot in the discovery sample in the *LEPR* locus at six months of age showing (a) the signal with lead SNP rs2767486 without conditioning, and (b) a putative second signal with lead SNP rs17127815 after conditioning on rs2767486.

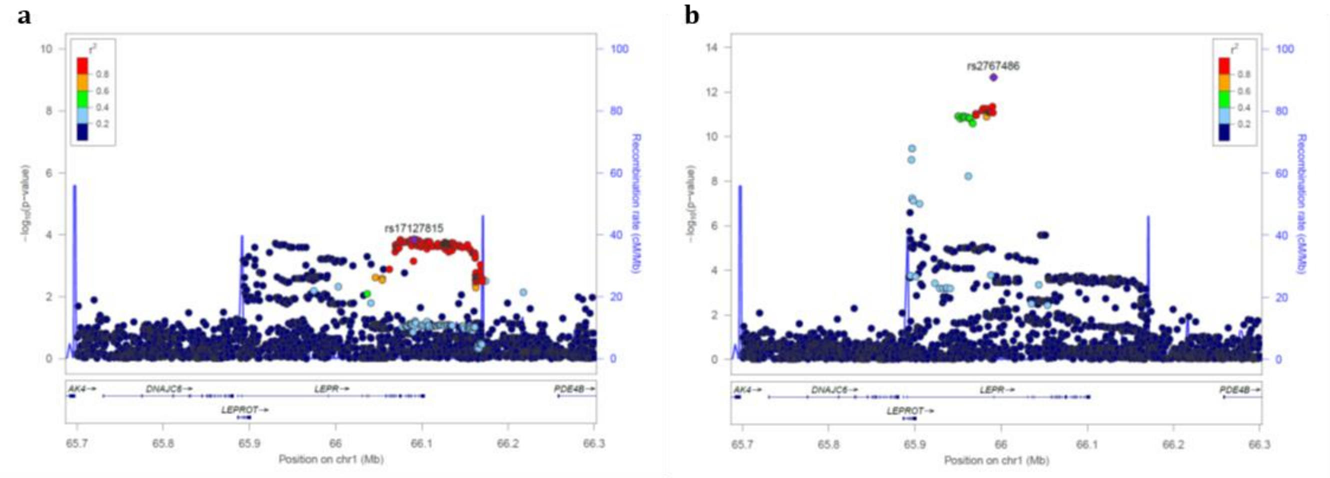

***Supplementary Figure 4***

**Conditioning on variants associated with related traits.** Regional association plot in the discovery sample in the *LEPR* locus at six months of age after conditioning on (a) rs2767485, the strongest pQTL for sOB-R-plasma levels in adults, and (b) rs11208659, associated with severe obesity. See main text for respective references.

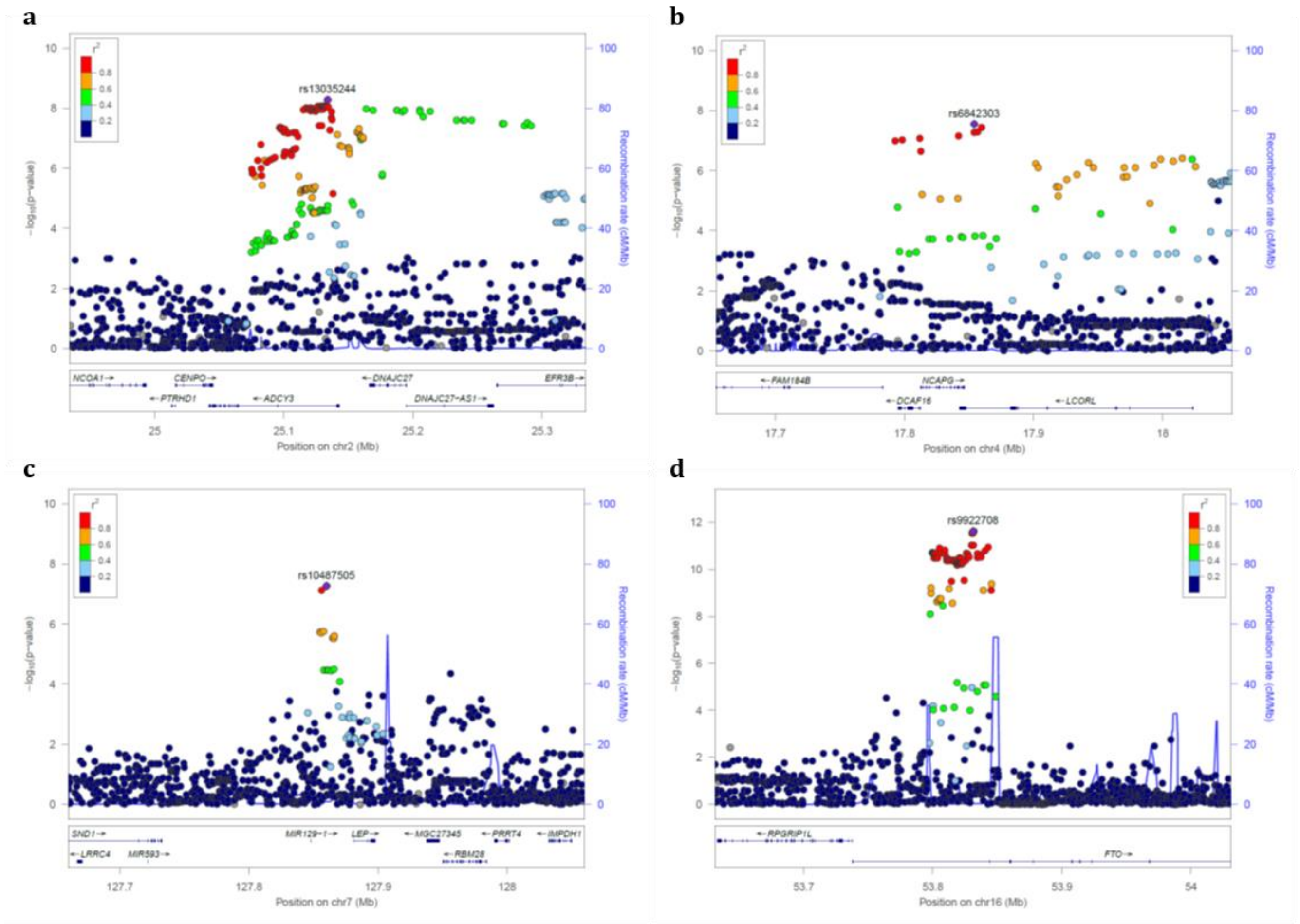

***Supplementary Figure 5***

**Association in the *ADCY3*, *LCORL*, *LEP*, and *FTO* loci.** Regional association plot in the discovery sample (a) rs13035244 in the *ADCY3* locus at 1 year, (b) rs6842303 in the *LCORL* locus at 1.5 years, (c) rs10487505 in the *LEP* locus at 1.5 years, and (d) rs9922708 in the *FTO* locus at 7 years.

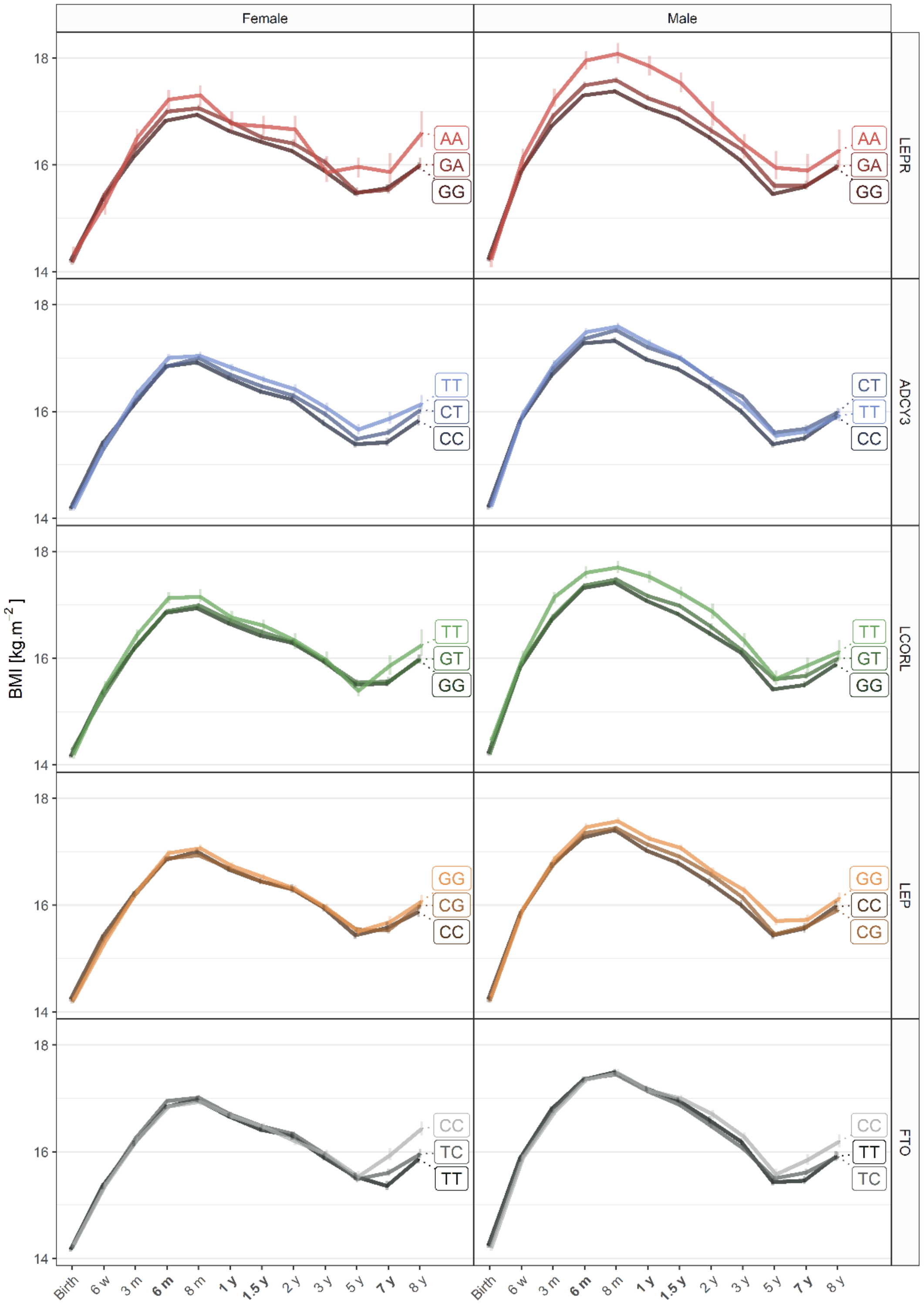

***Supplementary Figure 6***

**BMI trajectories stratified by the genotype of rs2767486, rs13035244, rs6842303, rs10487505, and rs9922708, lead SNPs of the *LEPR*, *ADCY3*, *LCORL*, *LEP*, and *FTO* loci, respectively.** The median of the BMI in $kg.m^{-2}$ at each time point for all samples (discovery + replication) stratified by genotype in dark, intermediate, and light colors for females (left) and males (right). Error bars represent 1 standard error of mean (SEM) on each side of the point.

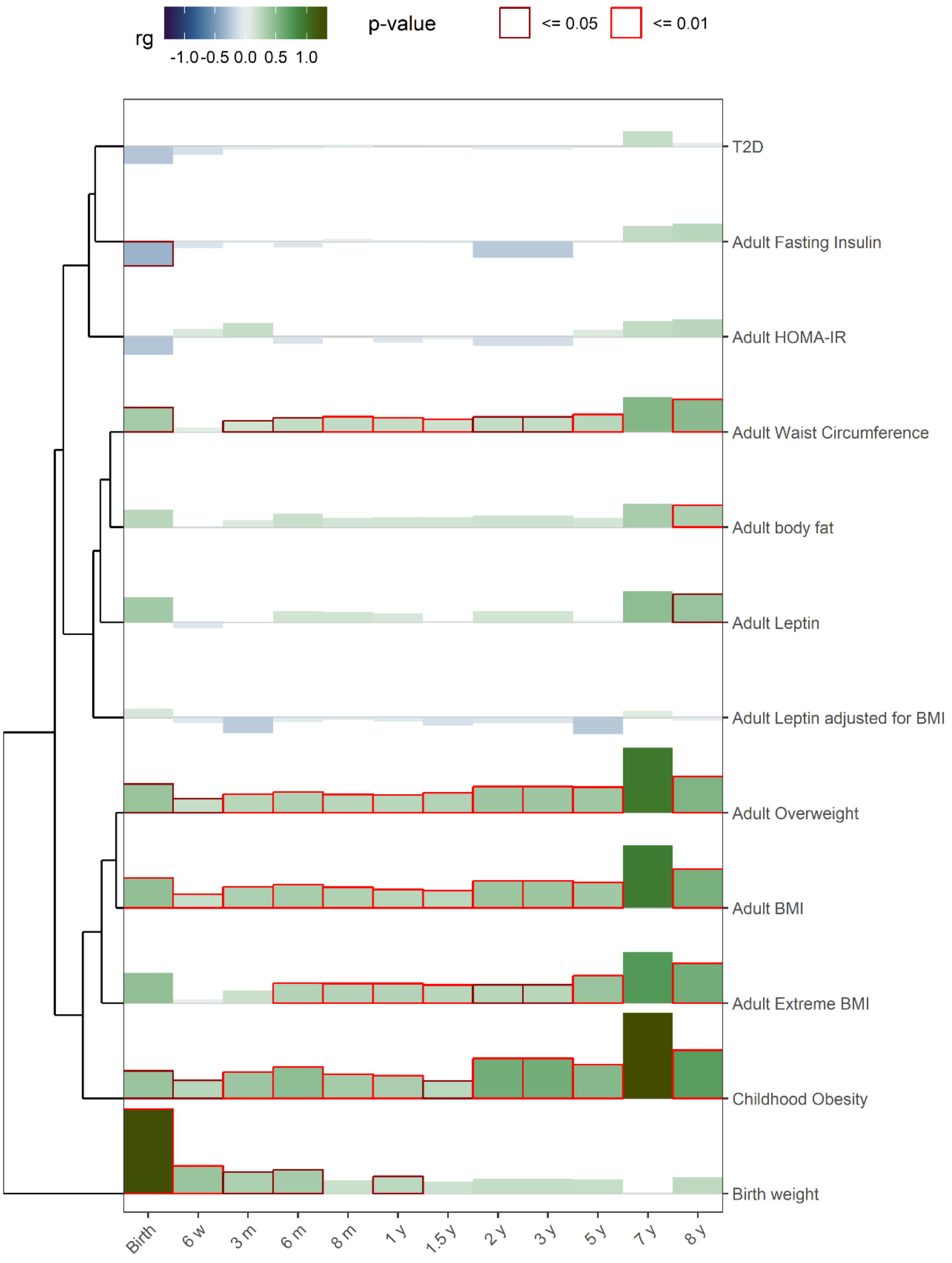

***Supplementary Figure 7***

**LD-score regression coefficient (rg) between BMI and related traits.** The LD-score regression coefficient ($r_g$) between our BMI association results at each age and related traits is plotted at each time point as column of height and color representative of $r_g$. Each column is outlined with a dark red and red square for p-values ≤0.05 and ≤0.01, respectively. The traits are ordered after hierarchical clustering as displayed with the dendrogram to the left: (i) Type 2 Diabetes, T2D, PMID 22885922; (ii) Fasting insulin main effect, PMID 22581228; (iii) homeostatic model assessment for insulin resistance, HOMA-IR, PMID 20081858; (iv) Waist Circumference, PMID 25673412; (v) Adult body fat, PMID 26833246; (vi) Leptin not adjusted for BMI, PMID 26833098; (vii) Leptin adjusted for BMI, PMID 26833098; (viii) Adult overweight, PMID 23563607; (ix) Adult BMI, PMID 20935630; (x) Extreme adult BMI, PMID 23563607; (xi) Childhood obesity, PMID 22484627; and (xii) Birth weight, PMID 23202124.

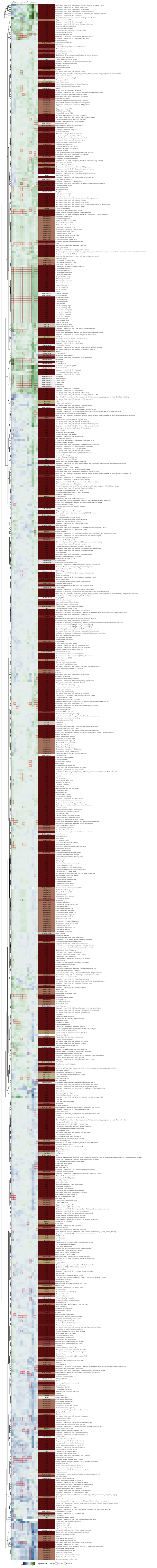

***Supplementary Figure 8***

**LD-score regression coefficient (rg) between BMI and other traits.** The LD-score regression coefficient ($r_g$) between our BMI association results at each age and other traits is plotted at each time point as heatmap of color representative of $r_g$. Each cell is outlined with a dark red and red square for p-values ≤0.05 and ≤0.01, respectively. The traits are ordered after hierarchical clustering as displayed with the dendrogram to the left.
